# CogStack - Experiences of Deploying Integrated Information Retrieval and Extraction Services in a Large National Health Service Foundation Trust Hospital

**DOI:** 10.1101/123299

**Authors:** Richard Jackson, Ismail Kartoglu, Asha Agrawal, Kenneth Lui, Honghan Wu, Tudor Groza, Angus Roberts, Genevieve Gorrell, Xingyi Song, Damian Lewsley, Doug Northwood, Amos Folarin, Clive Stringer, Robert Stewart, Richard Dobson

## Abstract

**Background:** Traditional health information systems are generally devised to support clinical data collection at the point of care. However, as the significance of the modern information economy expands in scope and permeates the healthcare domain, there is an increasing urgency for healthcare organisations to offer information systems that address the expectations of clinicians, researchers and the business intelligence community alike. Amongst other emergent requirements, the principal unmet need might be defined as the 3R principle (right data, right place, right time) to address deficiencies in organisational data flow while retaining the strict information governance policies that apply within the UK National Health Service (NHS). Here, we describe our work on creating and deploying a low cost structured and unstructured information retrieval and extraction architecture within King’s College Hospital, the management of governance concerns and the associated use cases and cost saving opportunities that such components present.

**Results:** To date, our CogStack architecture has processed over 300 million lines of clinical data, making it available for internal service improvement projects at King’s College London. On generated data designed to simulate real world clinical text, our de-identification algorithm achieved up to 94% precision and up to 96% recall.

**Conclusion:** We describe a toolkit which we feel is of huge value to the UK (and beyond) healthcare community. It is the only open source, easily deployable solution designed for the UK healthcare environment, in a landscape populated by expensive proprietary systems. Solutions such as these provide a crucial foundation for the genomic revolution in medicine.

## Background

Large healthcare organisations are often responsible for provisioning care in a wide range of medical specialties. It is not uncommon for a given speciality to make use of bespoke IT systems to support the specific requirements of clinicians at the point of care, such as imaging technologies, electronic prescribing and intensive care monitoring. This leads to a tendency for healthcare IT departments to support a large number of systems, which often suffer from integration issues, in the sense that there may not be a single interface that allows users to access data across all systems simultaneously. While there have been many attempts to standardise intra-system communication with the use of controlled languages and data schemas, such as HL7[1], the myriad of vendors, differential versioning of the standards and the ambiguity in the interpretation of the standards has caused such efforts to be only partially successful in practice[2, 3, 4]. This has lead to a high degree of heterogeneity in how information is managed within and between different NHS Trusts, which in turn has inflated the costs of creating suitable data management and analytics solutions, due to the investment required for successful implementation. For the end user, whether they be a clinician, a researcher or a business intelligence analyst, the implication is often described as a ‘needle in a haystack’ problem, owing to the complexity of how, where and why data is stored in a host of disparate sources. Without significant guidance from central hospital IT departments, many lay users of health information systems may not be aware of the logic of how data flows between them, and thus opportunities to use the organisation’s data to drive efficiency improvements are undermined.

The problem is further compounded by the nature of health data. In contrast to domains where structured data are captured in abundance (for example in e-commerce customer behaviour, retail loyalty card usage and financial trading patterns), all but a thin supernatant of clinical information are recorded as unstructured data in the form of the clinical narrative, via free text clinical notes, discharge summaries and referral letters[5, 6]. Since unstructured data are inherently more difficult to manage and query, this preference of clinicians manifests as a complication in how data can be provisioned between stakeholders effectively.

Information retrieval technologies have the stated aim of providing the ability to filter very large quantities of both structured and unstructured information and return relevant results at high speed. Due to their relatively straight-forward manner of ingesting data without a requirement to pre-define a schema, they have enjoyed a long history of success in almost every domain of information management, and are deployed in business critical environments such as enterprise document retrieval, bioinformatics, e-commerce and log management. Typically, they are provisioned through a simple, intuitive interface by which a user can query structured and unstructured data simultaneously, and rapidly refine their query to provide results relevant to their intent. This feature of query refinement through iteration is especially important in healthcare, given the nature of the medical ‘sub-language’, where concepts tend to be represented in clinical text with a high degree of assumed knowledge and a low level of verbosity[7, 8].

When correctly implemented in a healthcare organisation, such technologies are increasingly employed to overcome a range of data accessibility issues. We delineate these issues by what we refer to as the 3R principle:

### Right Data

With large amounts of data flowing through an organisation, often conflicting reports may occur. For example, two different diagnoses may be reported on two separate occasions. A third party who only has access to a partial view of the data would not be able to make a judgement on the current status of a particular patient. Therefore, maximising the recall (sensitivity) of an information retrieval system is essential to ensure data sufficiency for a question answering system. On the other hand, almost a decade of widespread EHR adoption has created a deluge of data in many progressive healthcare organisations - a trend which is certain to grow. A key consideration reflecting the usability of an information retrieval system is therefore also its ability to avoid false positives (precision, or positive predictive value) and not overburden the user with irrelevant results.

### Right Place

Many enterprise grade approaches to integration opt for data-warehousing methods to provide a single end point, often a SQL relational database, to offer an online analytical processing (OLAP) style capability. While the value of such approaches is well established, it is often restricted to users of the business intelligence community, and generally limited in its ability to effectively manage free text. This constraint therefore inhibits users elsewhere in the organisation, who may have simpler requirements regarding data use (for example, to find documents relating to patients in their care that contain certain keywords). In addition, the technical skills required to use OLAP resources effectively may concentrate in a relatively small number of individuals. Therefore, user-friendly solutions with a lower technical barrier for effective use will enable a degree of ‘self provisioned analytics’ and thus enjoy a wider uptake amongst employees.

### Right Time

Time based factors are often the difference between actionable and ‘stale’ information in clinical and business decision making. For instance, the opportunity to code clinical documents for repatriation may be lost if relevant data cannot be supplied to a code billing team within a commercial deadline. Similarly, if the data deluge negate the possibility of a human reading every document, there is potential to under-code the dispensation of high cost drugs and/or services. In the case of critical care, identifying antagonistic factors towards recovery at speed may help to deliver more favourable outcomes. The requirement to make data available throughout the organisation with as little latency as possible is critical to ensure its effective use.

### Information Governance

The aim of our project is to offer a general information retrieval system and OLAP analytics capability to meet the requirements of a large variety of use cases. However, in order to protect the rights of individuals as per the UK 1998 Data Protection Act, there are strict controls on how different types of data can be used for different purposes. From a technical perspective, this imposes limitations on how and where data can be provisioned and what transformations it must undergo. Generally speaking, the individuals within a given dataset may be classed as identifiable (no information is removed), pseudonymised (identifiers replaced by a pseudonym, enabling data linkage to other datasets), or anonymised (all identifiers removed, or data aggregated such that re-identification of individuals is nearly impossible). Each class of information removal represents different levels of risk regarding the secondary use of data. Although the details of the Act are complex, the practical applications in a clinical setting might be summarised in the following scenarios:

### Business Intelligence

Activities that utilise the data a Trust holds for the purposes of improving its operational efficiency. Here, named functions within the Trust may use identifiable data for a limited number of well defined purposes. For example, the Trusts clinical coding function has the remit to examine data generated in the course of a patient’s care, to ensure that delivered clinical services are accurately recorded and billed for. Alternatively, the Trust may use its data to meet its legal requirements to report figures to central government departments concerning the organisation’s performance or indicators of the nation’s health, such as cancer survival or diabetes rates.

### Service Improvement Activities

Under approval from the Trust’s appointed Caldicott guardian, Trust staff may access pseudonymised data in limited amounts in order to undertake internal research projects with the aim of improving the quality and/or efficiency by which a Trust delivers clinical services. The criteria for this generally requires that the affected patients will potentially directly benefit from the project outcomes. For example, this scenario might be invoked if a clinician is seeking to challenge current practices in service delivery, such as how the length of inpatient of hospital visits are predicted in order to reduce the number of staff hours invested in this task.

### Enclave Style Research Environment

An increasingly common method by which non-staff researchers are able to access clinical record data. Similar to service improvement activities, this method covers an expanded scope that enables clinical data to be used for research projects beyond direct patient benefit. Here, external parties may access pseudonymised and de-identified clinical data in limited quantities in highly secure environments under ethical agreements granted by UK Research Ethics Councils. Examples include Clinical Records Interactive Search[9] and Secure Anonymised Information Linkage Databank [10].

### Explicitly Obtained Consent

Perhaps the most common method of accessing clinical data for research is by explicitly obtaining consent from patients to use their identifiable data. This is also governed by Research Ethic Councils, and generally involves strict practices to guard against data breach. Although the most liberal in terms of how the data can be used (since patients are directly briefed as to the nature of the research and how their details will be used), the resource intensive means by which consent must be obtained generally creates a practical limit on the number of patients that can be included in such studies. In turn, this affects the type of study for which this approach is suitable.

## Implementation

Here, we describe our work on the CogStack architecture, an open source information retrieval and extraction architecture to provide an alternative to the UK healthcare community in a space traditionally occupied by commercial vendors. We describe its features and how it has been implemented within King’s College Hospital (KCH). We focus specifically on surfacing the deep data with the EHR for identification and recruitment of patients into the 100k Genomics England Project[11],for which the concept was funded and developed via NHS England Enablement Funding. Finally, we explore a vision of how such technology can be exploited for a range of use cases within the modern hospital environment.

### Previous Work

There are several reports of systems that offer information retrieval solutions directed at the challenges within the healthcare domain. Moen et al[12] proposed a variety for methods for selecting similar care episodes from other patients, given a particular case of interest. The NLP-Pier concept[13] combines an information retrieval and a biomedical entity information extraction system based around the popular open source project Elasticsearch. In the UK, comparable projects that make use of information retrieval systems include the CRIS[14] project, which uses the commercial FAST search engine and a custom text de-identification algorithm to make clinical notes from mental health patients available for research.

### Cognition

In addition, the open source Cognition platform[15, 16] is a vertically and horizontally scalable application that retrieves binary encoded documents and plain text from a relational database, and optionally de-identifies personal identifiers (for example, patient names, addresses and phone numbers) in text.

During routine clinical administrative activity, PHIs are often routinely collected as semi-structured data during the course of a patients care (for example, patient and carer names, addresses, NHS numbers and dates of birth). Such information is a valuable source of data for de-identification methods, as it offers highly precise information about the nature of the text strings that should be removed. However, in natural language, PHIs are often written in a variety of formats, requiring that high accuracy approaches have a greater flexibility that can be achieved by simple direct string matching. For instance, an address written “Institute of Psychiatry, Psychology and Neuroscience, 16 De Crespigne Park, SE5 8AF” might be shortened to “Institute of Psychiatry, 16 De Crespigne Park SE5 8AF”. Similarly, PHIs in natural language documents may contain spelling mistakes or additional punctuation tokens. To achieve flexibility, the effectiveness of rules based approaches has been demonstrated elsewhere[17]. The Cognition de-identification algorithms, which are used in this work, are designed to take into account misspellings, tolerance for missing/redundant information, and word order without the need for manual rule crafting nor construction of labelled datasets for machine learning approaches, which are known to be an expensive process[18]. Cognition applies a “sliding window” approach to detect the regions of text where patient identifiers are mentioned. During the processing of a document, the patient specific PHIs are retrieved from semi-structured fields in a database, and the Levenstein edit distance is calculated for each PHI token at every character offset available in the document. If the Levenstein distance is above a configurable threshold, the offsets of the match are masked. This allows for an efficient method of removing PHIs in a document, even if they are misspelt in the document or source inputs.

The de-identified output text from Cognition contains meta-data related to patients and the document such as a hash code of a combination of the patient’s identifiers and document date, which are useful for version control. The output text may also be output to a relational database or Elasticsearch index, to be used by downstream services such as the Kibana web interface, or natural language processing applications. Cognition uses the Apache Tika library for converting common document formats such as Microsoft Word, PDFs, Excel etc. into text and further applies Optical Character Recognition to scanned documents that are only available in image formats (including scanned PDFs) using the Tesseract library. Cognition handles horizontal scaling by using a HTTP-based coordinator-client approach where a coordinator assigns work coordinates to the clients.

### The CogStack Architecture

CogStack is a set of open source and open core services, co-ordinated by a batch processing framework that builds on the concepts of the Cognition platform by offering additional interfaces for NHS systems and NLP technologies. Out-of-the-box open source components were selected from a variety of successful open source and freely licensed projects. The services can be deployed using the Docker containerisation technology, to maximise ease of deployment.

The overall goal of the architecture is to undertake a series of configurable transformations of clinical data housed in relational databases and to load the transformed data into an Elasticsearch information retrieval engine (otherwise known as a search engine - described below), whereupon the 3R principles can be more readily addressed than via direct communication with the untransformed source databases alone. Each transformation is highly configurable, in accordance with the desired use case of the end product. For example, it is not necessary (or even desirable) to de-identify data for business intelligence use cases, and thus this can be disabled. Similarly, not all use cases will require computationally expensive entity extraction NLP processes. The rationale for the choice of components is described below, while the flow of data and transformations in the CogStack architecture is described in figure 1.

**Figure 1.**
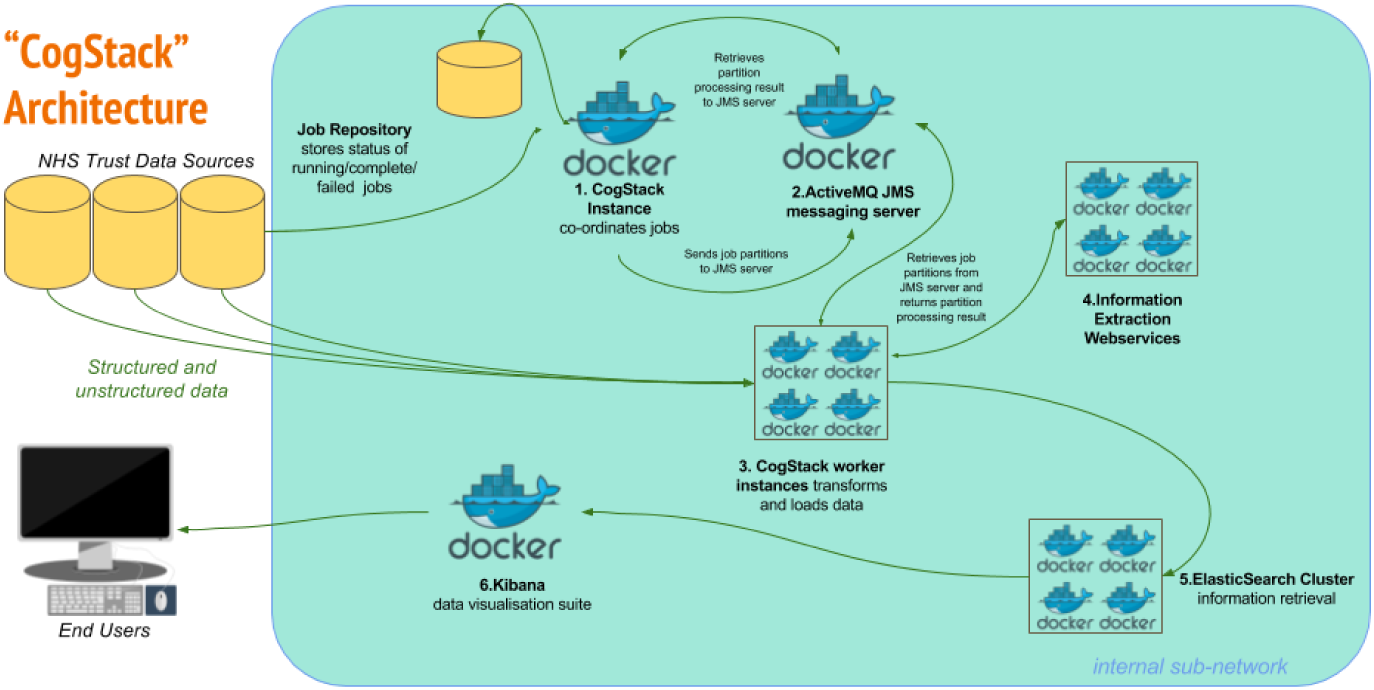
CogStack Architecture and Dataflow. All components can be deployed via the Docker containerisation software. **1. New job** execution Master instance of CogStack identifies new data in Trust Data Sources at intermittent intervals.**2. Partitioning** The job is partitioned into a user definable number of work units. **3a**. **Derive the freetext content** Extract plain and/or formatted text from common proprietary document binary formats (performing OCR where necessary), using the Tika Library to enable the downstream processing of high value unstructured data elements.**3b. Supplement the text content with meta-data** Filter and de-normalise a subset of the structured clinical data to provide a patient orientated, transparent representation of high value metadata concepts. For example, this might include calculated fields to represent patient age at document date, first part of postcode and ethnicity and lab results.**3c. De-identification** Transform the resulting text documents into de-identified text documents, by masking personal health identifiers via the use of the Cognition de-identification algorithms. This is necessary to address governance concerns associated with the secondary use of patient data. Identifiers in structured data can be excluded via SQL query, according to business requirements. **4. Information Extraction** Apply generic clinical IE pipelines to derive additional structured data from free text and supplement the quantity of available structured data at the point of query. **5. Indexing** Build a JSON object from the resulting structured and unstructured data, which can then be readily be indexed into an Elasticsearch cluster. **6**. **Visualisation** The Kibana suite provides a range of attractive options for viewing, aggregating and dash-boarding the loaded data

### Handling Text and Other Unstructured Data

During a patient’s course of treatment, a large number of documents such as referral letters and discharge summaries tend to be generated via word processing applications, predominently Microsoft Word. In addition, such documents may undergo further manipulations, such as PDF conversion and printing and rescanning as an image before they reach their final storage location, usually a relational database. Such manipulations represent complications for search and NLP applications, as the valuable electronic free text content may be ‘locked’ inside proprietary file formats, or even lost during the conversion to an image format. The Apache Tika library[19] provides the capability to extract electronic text from a wide variety of file formats, and (in combination with the open source Tesseract Optical Character Recognition (OCR) tool[20]), recover images of text back into character electronic format. At the time of writing, Tika does not provide the capability to perform OCR on PDFs containing images. To this end, we enhance Tika with a custom PDF parser class, additionally making use of the ImageMagick tool in order to generate the required inputs for use with Tesseract.

### Biomedical Entity Extraction, Bio-YODIE and BioLark

Implementing an information retrieval system over clinical records represents a high return on investment by lowering the barrier to large scale data access in line with the 3R principle. However, the limitations of information retrieval are well recognised in terms of its ability to deal with ambiguity, different word senses, negation and other factors that are likely to produce an irrelevant or imprecise result. In order to provide a higher granularity of data at the point of search, it is necessary to implement information extraction (IE) techniques to enhance text elements with meta-data. To this end, the CogStack architecture offers two third party pipelines, with the capability to extend the system with additional pipelines via webservices.

First, Bio-YODIE is a clinical information extaction system designed for use with UK clinical records. It development was necessitated in response to the widely recognised generalisability issues of English language clinical NLP systems, which have historically arisen in the United States[21, 22]. Bio-YODIE is designed to extract a subset of Unified Medical Language System[23] concepts in free text. This subset is selected for their high business value, and includes entities such as drug names (both brand and generic), disease names and procedures. The full UMLS system is not used to reduce noise and retain scalability.

Bio-YODIE has been evaluated against two corpora; the MIMIC II corpus [24, 25] and a new corpus created using patient records at the South London and Maudsley NHS Foundation Trust. In the latter, 201 documents have been triple-annotated by medical experts, achieving a three way interannotator agreement of 0.747. The corpus is confidential; however annotator guidelines are available for public review[26]. Bio-YODIE achieves an accuracy of 0.926 on the task of correctly linking to UMLS concepts on the SLAM corpus, 0.842 on the MIMIC 2013 test set and 0.827 on the MIMIC 2014 test set. A separate evaluation of NER performance (finding the right parts of the text, rather than as above, disambiguating them correctly given that the span has already been located) shows that Bio-YODIE achieves an F1 of 0.751 on perfect span matches (0.823 when concepts with any degree of overlap are also counted) on the SLAM corpus; however when only correct types are counted, this falls to 0.523 (0.564). NER performance was not evaluated on the MIMIC corpus because this corpus is not fully NER-annotated. In a comparative evaluation (forthcoming), Bio-YODIE and MetaMapLite offered similar advantages over the competitors considered in terms of accuracy, speed and stability; however, Bio-YODIE also offers the possibility to include prior probabilities from corpus data, resulting in a substantial improvement in disambiguation accuracy. For this reason, Bio-YODIE was chosen. Bio-YODIE is dual licensed under GNU Affero and commercial options.

Second, Bio-lark encodes clinical text with Human Phenotype Ontology[27] concepts - the principle ontology for phenotyping patients in the 100K Genomics England Project. Negation detection for HPO terms is provided by the NegEx algorithm[28]. An evaluation of the system over a Pubmed corpus is described in [29]. Here, Biolark achieved an F1 score of 0.95 over a test set of 1 933 instances, corresponding to 460 unique HPO concepts. Bio-lark is available under an academic license.

The outputs of the NLP processes are captured as JSON objects and indexed using the ‘nested’ type of Elasticsearch. In doing so, it is possible to query unstructured data as though it were structured, although the accuracy will vary greatly depending on a multitude of factors.

### Text De-identification Performance

Different use cases for Trust data have different governance requirements. The requirements for the anonymisation and pseudonymisation has been the subject of national and international working groups[30, 31, 32]. The process of masking Protected Health Identifiers (PHIs) in clinical free text remains an active area of research from both a governance and NLP perspective. The Informatics for Integrating Biology and the Bedside (I2B2) organisation regularly organises open challenges for NLP researchers to examine the state of the art in text de-identification technology, by providing corpora of PHI annotated clinical text for international researchers to experiment with[33]. Such efforts have undoubtedly yielded significant advances in the field, to the extent that the performance of hybrid knowledge driven and machine learning methods equals that of human annotated documents in controlled test environments. Nevertheless, there remain outstanding tasks to ensure that such approaches are generalisable across different languages, dialects, specialities and hospital systems.

Due to strict data protection laws, it is generally not possible for researchers to access clinical text containing identifiable information. Therefore, validating the Cognition de-identification algorithms poses certain challenges. While certain domain corpora are available via activities such as I2B2 described above, these are not representative of UK clinical data. Therefore, we created a simulated dataset to explore the performance on registered company address entities harvested from public records. We devised a series of string mutator methods to attempt to recreate a variety of likely scenarios that would cause named entities to vary between two sources. These mutation methods were designed to represent real world events that might cause clinical document PHIs to not match those entered via an administrative process, and thus limit the effectiveness of exact string matching. We decided to focus on address named entities only, as these tend to offer the greatest scope for variation, compared with first/last name, telephone number and NHS number PHIs.

We explored four types of mutation method. First, keyboard typographic errors using prior probabilities of frequently mistyped keys, at a per character error rate of 3%, 10% and 20% (for example, ‘100 Meadow Street’ to ‘100 Meagow Streat’. Second, substituting full address tokens to common abbreviations and vice versa (for example, ‘Road’ to ‘Rd’ and ‘St’ to ‘Street’) at a per token rate of 100% (i.e. any detected possible address substitutions were replaced). Third, an address token truncator, which removes tokens from the end of an address. The purpose of this is to replicate the observation that in some cases, full addresses (often supplemental address lines) are not recorded. For instance, ‘100 Meadow Street, Barkingford, Greater London, London’ may be shortened to simply ‘100 Meadow Street’. We specified a token removal rate of 100%, with a minimum address length of three tokens. Finally, the most convoluted mutator we implemented was designed to mimic the effects of poor quality OCR. This mutator includes the effects of the character substitution mutator, with the additional possibility of inserting whitespace characters at random intervals within tokens. We tested this mutator with a per character substitution rate of 3%, 10% and 20%, and a per character whitespace insertion rate of 3%, 10% and 20%

The mutated address strings were then wrapped in ‘Lorem Ipsum’ style generated text to simulate surrounding language. We generated 1000 test documents under a variety of degrees of PHI mutation and report precision and recall statistics for per token masking.

### Scalability and Database Synchronisation

Scalability is achieved using the remote partitioning concept. Here, a unit of work is defined as a job (for example, to process 10 000 rows of new/updated data since the last job was executed). A master process partitions this job into a configurable number of smaller work units. These partitions and other job metadata are stored in a job repository and then sent as a message to a Java Messaging Service (JMS) server. These are then picked up by multiple worker processes operating on local or remote servers. Upon the arrival of a partition, each worker will begin to execute the work described within the message. Upon completion, the worker processes will inform the master process (again via JMS) about the status of the partition. If all partitions are successful, the job will be marked as complete, and a new job will start to process any new data generated by business activities during the processing of the previous job. Via this mechanism, a degree of ‘near real-time’ synchronisation with the source databases are achieved, although in practice it is constrained by available hardware, database configuration and network speed.

### Elasticsearch and Kibana

Following the data transformation steps, the data is loaded into Elasticsearch, a popular open source search and analytics engine developed by Elastic.co. The non-transactional, NoSQL data model used by Elasticsearch enables the ingestion of large quantities of data at high speed, making it rapidly available for querying. Elasticsearch was chosen as it offers a number of advantages over traditional relational databases, predominantly concerning it’s advanced capabilities to construct complex queries over structured and unstructured data simultaneously. In addition, the NoSQL data model it supports enables schema free loading of data (in the sense that there is no need to predefine the structure of data before it is loaded). This is particularly advantageous given the myriad of different database systems supported within a typical NHS Trust, as the technical debt incurred by connecting new data sources to the engine is greatly reduced. As an analytics engine, Elasticsearch allows common and complex aggregations to be performed at speed. Finally, Elasticsearch offers a Representational State Transfer (REST) web service, which cab be flexibly leveraged to allow external applications and services to retrieve data using the HTTP protocol.

For the end user experience, the open source Kibana data visualisation application (also by Elastic.co) is specifically designed to interact with Elasticsearch, and offers document visualisation, text highlighting and dashboarding capabilities. Via Kibana, non-technical users are able to search document text and structured meta-data in much the same way as one would use an e-commerce website. A screen shot of the Kibana interface is provided in figure 2.

**Figure 2.**
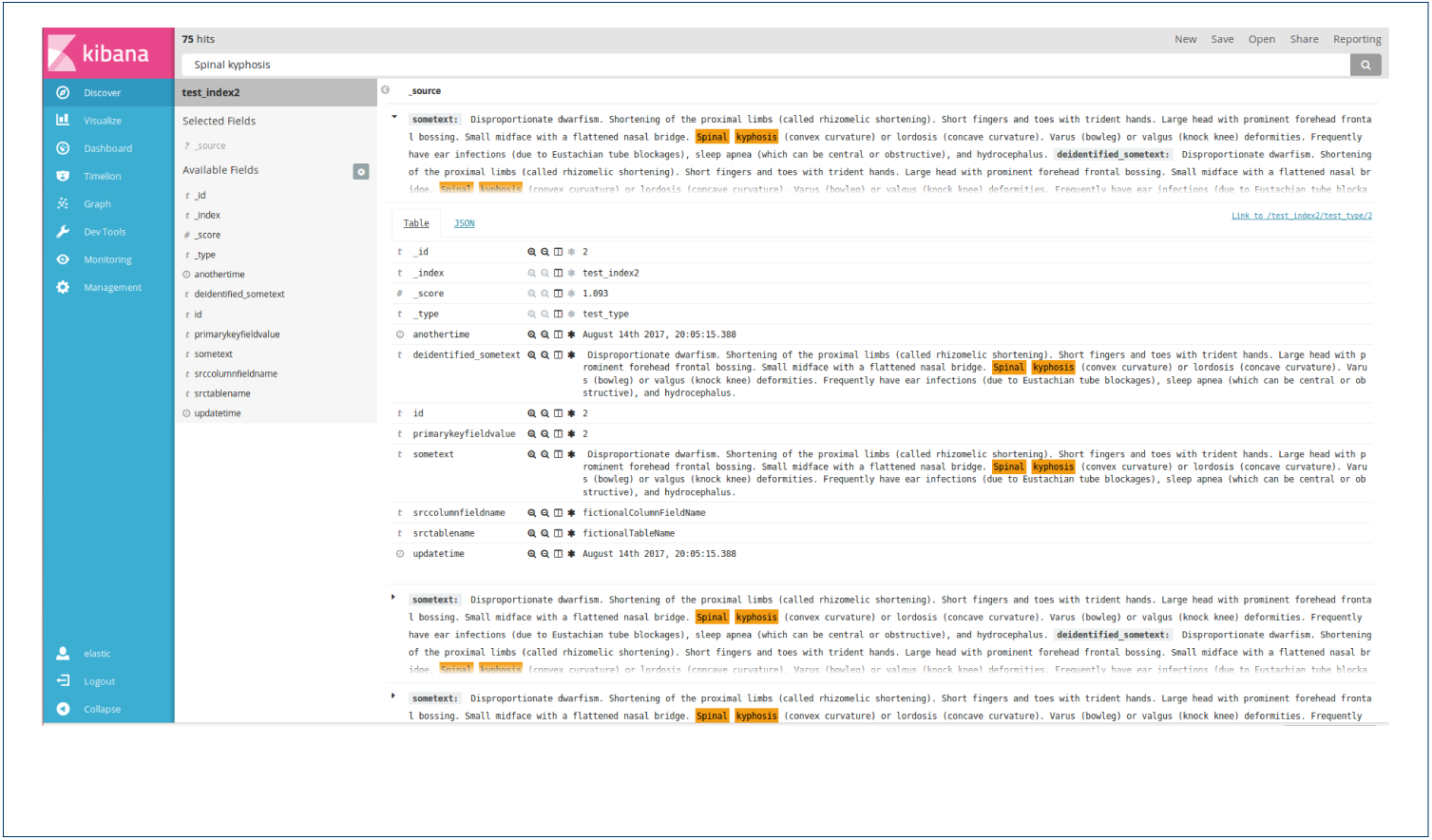
Kibana interface loaded with pseudo-data.

### Security and Information Governance

Due to the sensitive nature of the clinical data, access is administered via a system manager in line with the information governance scenarios described above. Technical considerations are managed via commercial grade security provided by Elasticsearch plugins, offering active directory/LDAP/HTTP user authentication control, user access logging for audit, per index access restrictions with optional document/field level access restrictions and private certificate authority SSL encryption to protect in-flight data.

## Results/Discussion

### Data Model

As of December 2016, we have used the CogStack architecture to process approximately 300 million rows of clinical data from KCH databases. This data has been organised into identifiable and de-identified indexes for business intelligence and service improvement concerns, trial recruitment and tailored care use cases.

Each index is centred around a high value concept-

### Observations

Clinical notes taken during patient/doctor interactions (24 991 406 rows)

### Basic Observations

Test results from pathology systems and short notes (248 028 823 rows)

### Orders

Prescribing information (66 838 164 rows)

### Documents

Binary documents generated by inter and intra Trust communication, comprising 8 736 295 rows. Of these, 4 505 750 (52%) resulted from MS Office, 3 479 583 (40%) were PDFs and 340 764 (4%)required OCR

#### Demographics of the Acute Patient Population at King’s College Hospital

A short demographic breakdown of patients across all years is given in table 1. Top level ICD-10 groups, as assigned by clinical coders are presented in table 2.

**Table 1.** Patient demographics, King’s College Hospital 2004-2016

|  | Count | % |
| --- | --- | --- |
| Age (years) |  |  |
| ≤ 20 | 435 796 | 14.80 |
| 21–40 | 811 865 | 27.57 |
| 41–60 | 876 467 | 29.77 |
| 61–80 | 490 153 | 16.65 |
| ≥ 80 | 326 453 | 11.09 |
| Unknown | 3 792 | 0.13 |
| Gender |  |  |
| Male | 1 369 074 | 46.50 |
| Female | 1 571 717 | 53.38 |
| Indeterminate | 550 | 0.02 |
| Unknown | 3 185 | 0.11 |
| Race (Self Assigned) |  |  |
| Asian or Asian British | 95 682 | 3.25 |
| Black or Black British | 326 618 | 11.09 |
| Mixed | 59 214 | 2.01 |
| Not specified | 1 506 703 | 51.17 |
| Other | 9 7277 | 3.30 |
| White | 859 032 | 29.17 |

**Table 2.** ICD10 Code assignment by clinical coders at King’s College Hospital

| Group | Unique Patient Count |
| --- | --- |
| I Certain infectious and parasitic diseases | 171 988 |
| II Neoplasms | 259 975 |
| III Diseases of the blood and blood-forming organs and certain disorders involving the immune mechanism | 72 939 |
| IV Endocrine, nutritional and metabolic diseases | 272 317 |
| IX Diseases of the circulatory system | 504 581 |
| V Mental and behavioural disorders | 706 990 |
| VI Diseases of the nervous system | 179 710 |
| VII Diseases of the eye and adnexa | 183 841 |
| VIII Diseases of the ear and mastoid process | 13 416 |
| X Diseases of the respiratory system | 242 282 |
| XI Diseases of the digestive system | 598 165 |
| XII Diseases of the skin and subcutaneous tissue | 131 227 |
| XIII Diseases of the musculoskeletal system and connective tissue | 343 803 |
| XIV Diseases of the genitourinary system | 212 198 |
| XIX Injury, poisoning and certain other consequences of external causes | 351 608 |
| XV Pregnancy, childbirth and the puerperium | 327 111 |
| XVI Certain conditions originating in the perinatal period | 78 541 |
| XVII Congenital malformations, deformations and chromosomal abnormalities | 104 242 |
| XVIII Symptoms, signs and abnormal clinical and laboratory findings, not elsewhere classified | 513 384 |
| XX External causes of morbidity and mortality | 650 984 |
| XXI Factors influencing health status and contact with health services | 1 520 107 |
| XXII Codes for special purposes | 385 417 |

#### Text de-identification validation

The results of our four methods to simulate PHI input errors for 1 000 addresses are given in table 3. Because of the use of a random number generator to determine when string manipulations should occur, the total number of PHI tokens varies slightly between executions. For each test, approximately 8 500 pseudo-PHI address tokens were generated. For our character substitution mutator, precision ranged from 93.9% at a 3% substitution rate to 96.3% at a 20% substitution rate. Recall ranged from 95.5% at a 3% substitution rate to 82.0% at a 20% substitution rate. Performance over address aliasing achieved 94.4% precision and 94.8% recall. For token removal, precision was calculated at 96.6% and recall 92.1%. Performance on simulated OCR documents performed the least well, with precision at 98.2% and recall at 84.5% at a 3% character substitution rate and 3% white space insertion rate. At 20% charater substition rate and 20% white spcae insertion rate, precision was 92.3% and recall was 11.0%.

**Table 3.**
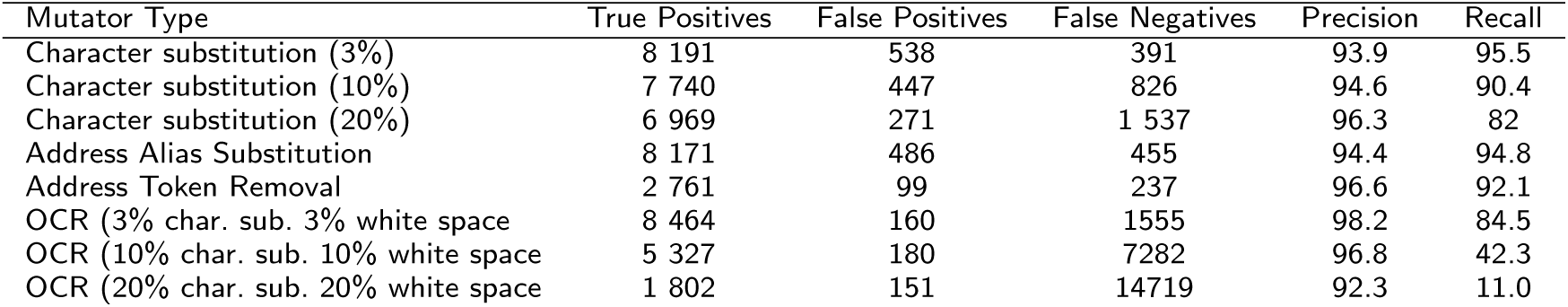
Performance of de-identification on simulated data

| Mutator Type | True Positives | False Positives | False Negatives | Precision | Recall |
| --- | --- | --- | --- | --- | --- |
| Character substitution (3%) | 8 191 | 538 | 391 | 93.9 | 95.5 |
| Character substitution (10%) | 7 740 | 447 | 826 | 94.6 | 90.4 |
| Character substitution (20%) | 6 969 | 271 | 1 537 | 96.3 | 82 |
| Address Alias Substitution | 8 171 | 486 | 455 | 94.4 | 94.8 |
| Address Token Removal | 2 761 | 99 | 237 | 96.6 | 92.1 |
| OCR (3% char. sub. 3% white space) | 8 464 | 160 | 1555 | 98.2 | 84.5 |
| OCR (10% char. sub. 10% white space) | 5 327 | 180 | 7282 | 96.8 | 42.3 |
| OCR (20% char. sub. 20% white space) | 1 802 | 151 | 14719 | 92.3 | 11.0 |

## Discussion

Our CogStack software arose out of a requirement from the 100 000 Genome project (100KGP) to find a low cost solution for providing relevant clinical data to the programme amongst the large volumes of disparate data sources within KCH. In response to this challenge, we have produced an open source integrated document retrieval and information extraction, to solve a variety of typical issues associated with analytics within an NHS environment. We chose a range of components on the basis of ubiquity, robustness, commercially friendly licensing and price to offer a viable alternative to commercial solutions. Beyond the initial scope of the 100KGP, our CogStack architecture has enabled us to transform and ingest a large volume of clinical data in a fashion consistent with the requirements for data reuse in business intelligence, service improvement and research.

### Case Study: Patient Recruitment into the 100 000 Genomes England Project

The 100 000 Genomes project is the largest human sequencing project in the world. It is a UK initiative to sequence 100 000 genomes from individuals suffering from various cancers and rare diseases, with the intent of developing a genomic medicine capability for the NHS. This will create new diagnostic criteria for patients, and contribute to research for new treatments and cures. While the ambitions are high, the logistical and technical challenges of delivering such a capability within routine care are substantial. Two areas of particular difficulty have been identified at KCH.

The first challenge is to find and contact eligible patients for recruitment. Genomics Medicine Centres around the UK (such as KCH) are responsible for the recruitment and data collection of patients into the project, using the various inclusion criteria specified by the co-ordinating body, Genomics England. One of the principal use cases for the CogStack architecture has been to offer the means to rapidly develop search criteria such that appropriate individuals are identified.

As noted by Moen et al[12], quantitatively validating the quality of results produced by an information retrieval system is a complex task, as identifying the relevance of results is often highly context specific. However, subjective reports of users of the system suggest that project staff are able to work with clinical care teams to navigate large quantities of structured and unstructured data, to find information required validate putative cases for recruitment and approach patients in their normal course of care. For instance, the system allows researchers to quickly assess which patients have records that contain pertinent keywords and/or UMLS concepts, a process that would have previously required significant technical skill, direct knowledge of patient cases or manual data trawling.

The second challenge posed is to subsequently surface the deep phenotype data from recruited patients. A requirement for acceptance into the 100KGP is the completion of an extensive patient phenotypic data model by the recruiting Genomic Medical Centre. Such data may be held in disparate systems, complicating its extraction. Similarly to the recruitment challenge, collating data is substantially easier if held in a single source with extensive search functionality. In addition, the added value of IE approaches to resolve relevance challenges such as word sense disambiguation and negation offer further options for data retrieval. The technologies that make up the CogStack architecture enables members of the 100KGP team to rapidly scour individual patient records, regardless of size, and efficiently extract the required information.

### Other use cases

While CogStack was built in response to the requirements of the 100k Genomics England project, its potential for a large number of other use cases was quickly realised. For instance, as previously described, clinical coding is the activity of hand curating clinical documents to identify the exercise of care activities, such as the prescription of drugs. Clinical coding is an important activity in acute care Trusts, as its efficiency affects the Trusts reimbursement from central government for care dispensed. The modern propensity to record and store large amounts of clinical and administrative data has created new challenges for clinical coders, owing to the increasingly unfavourable ratio of coding capacity to volume of data. Information retrieval and extraction technologies offer the potential for a substantial return on investment enabling clinical coders to navigate the data more efficiently. Such a capability is especially valuable in complex cases, where co-morbidity factors hidden amongst a mass of unstructured data can have a substantial impact in the accurate assessment of the cost of patient care.

In addition, one of the most useful tasks in an organisation with complex data flows is to be able to offer near real-time alerting capability. The commercial ‘Alerting’ plugin for Elasticsearch offers an easily configurable solution to send messages to a variety of endpoints, such as email addresses, REST webservices and enterprise communication software such as Slack and Hipchat. In a clinical setting, alerting clinical teams to events outside of their immediate jurisdiction may offer new opportunities for intervention. Within KCH, such capabilities are currently being explored in the following scenarios: 1) abnormal creatinine and CCP antibody levels to detect adverse reactions to methotrexate and pre-clinical rheumatoid arthritis respectively, to hasten communication between the Rheumatology and Pathology Departments 2) identification of previous evidence of adverse reactions such as rash in response to Sulfasalazine treatment (especially in emergency contexts) 3) monitoring for drug administration delays on wards 4) alerting of anti-coagulant team for patients being discharged on anti-coagulation therapy and 5) alerting of clinical intervention team if a high National Early Warning Score is detected. Presumably, such a list represents only a fraction of the scenarios that would benefit from the 3R principle. Pending further development and successful trials, a future goal will be to explore additional alerting scenarios.

### Additional Implementation Issues/Limitations

The secondary reuse of EHRs is complicated by several factors. Fundamentally, the clinotype and phenotype are related but different concepts in our semantics for heath datasets. The sufficiency and robustness of the clinical record is often called into question as a source of secondary research data[34, 35, 36]. For instance, our current deployment of CogStack at KCH does not have access to primary care data, and therefore cannot be said to offer complete patient profiles. Similarly, no effort has been made at this time to address the challenges of linking datasets across different secondary care organisations.

EHR data is predominantly used for front line recording and communication within care units. Missing data, or inconsistencies can be resolved if and when they become relevant to direct care by patient/care unit interaction. Such error correction routines are not possible in secondary use scenarios whereby corrective intervention is not feasible. The heterogeneous landscape of systems, data owners and APIs that are synonymous with IT infrastructure in large organisations are likely to compound the problem. Recording the same (or related) information in multiple systems, increases the likelihood of conflicts.

Governance, security and process issues require significant consideration in the development of standard operating procedures. It is likely that many Trusts have procedures in place to manage business intelligence, service improvement and research project with explicitly obtained consent. However, some of the most forward thinking opportunities for analytics require access to data at a scale where explicitly obtained consent is not feasible. Such activities likely require the use external resource and expertise, as has been the doctrine behind the establishment of NHS/University collaborations in the form of National Institute for Health Research Biomedical Research Centres. Few Trusts have the facilities to offer enclave style research environments to external researchers, for example in the form of the aforementioned CRIS and SAIL security models. This creates a significant limitation in the potential for localised secondary EHR use outside of such institutions, and discussions to address such issues continue to take place at the national level. Progress in this area is likely to take the form of substantial patient engagement activities to ensure the retention of public trust, and the development of pioneering models of consent such as Consent for Contact [37].

One particular factor of concern when managing unstructured data is the quality of OCR performance. Although only 4% of binary documents required OCR at KCH, our subjective assessment of the Tesseract library suggests OCR performance varies greatly in line with the quality of input. Good performance was observed when OCR was attempted on clean, printed black and white documents that were carefully aligned to scanner boarders. Deviations from these factors resulted in a rapid decline in OCR performance.

Regarding Information Extraction approaches, our efforts here offer Bio-lark and Bio-YODIE ‘out-of-the-box’ as a means to demonstrate compatibility with the CogStack concept. However, the necessity for domain adaption to new corpora of clinical text is well established[38, 39]. Future work will look the information extraction performance and ease of domain adaptation of these technologies to the KCH corpora.

The de-identification algorithms we make use of are deterministic string matching method based upon the same principles described in[17]. Although we were unable to validate the performance on real clinical data at this time, we would expect recall metric to be approximately the same.

Because of our access limitations to identifiable clinical data, we are hesitant to make broad comparisons with other methods in this area. We would have liked to compare performance across a range of algorithms, such as those proposed in the I2B2 2014 task for text de-identification. However, it should be noted that the majority of these algorithms are not available in the public domain. In addition, we note that such algorithms are designed for US style identifiers rather than UK ones, therefore requiring some form of domain adaptation for appropriate use. Regardless, our experiences of automated de-identification techniques suggest that appropriate ethical use should involve extensive internal validation on a per-dataset basis, before such data is deemed suitably transformed for further use cases.

Our testing of the approach in a simulated environment suggests reasonable performance of the de-identification algorithms to many forms of string perturbation, with the most noticeable drops in performance occurring with our ‘poor OCR’ simulations. It should be noted that at the higher grades of OCR error, documents became increasingly illegible, suggesting that PHIs may not be interpretable to human observers.

One particular dependency of the de-identification algorithm is that it requires PHIs to exist as structured or semi-structured fields in a database, which may make it unsuitable for some types of EHR data. Many other forms of PHI masker do not have this requirement[40]. However, due to the nature of its workings, it can synergistically be combined with other de-identification approaches.

Regarding resource allocation during the progress of the project, the most significant deployment cost arose from the need for the implementation team to understand the complex landscape of modern and legacy systems in place inside the Trust. For instance, these commonly took the form of certain services being unavailable at certain times, or restrictions on the load that could be placed on certain services to prevent interference with the day-to-day running of front line services. In such cases, it was necessary to retain flexibility with regard to requirements, in keeping with common agile management paradigms.

## Conclusion

Our CogStack software arose out of a requirement to build an integrated document retrieval and information extraction system for a large UK NHS Trust. Our experiences have led us to identify a variety of typical issues associated with the development of local analytics environments within the NHS, broadly encapsulated as what we define as the 3Rs of right data, right place and right time. We have released our software components under permissive licensing arrangements in the hope that other NHS Trusts might benefit from our findings.

### Availability and requirements

Project name: CogStack Project home page: The code, documentation, string mutator classes and example configurations for CogStack are available at https://github.com/RichJackson/cogstack/ Operating system(s): JVM based - The codebase should work on Windows and Linux systems, although Linux systems are recommended for docker style deployment Programming Language: Java, Groovy, Spring Batch Framework Other requirements: Java 1.8 or higher License: Apache 2.0 Any restrictions to use by non-academics: Please check with Angus Roberts and Tudor Groza before using the Bio-Yodie and Biolark components respectively Note: Instructions on how to reproduce the simulated data and results described here are included in the associated documentation, available at the same location.

For information governance reasons, no clinical record data can be provided with this research.

### List of abbreviations

NHS: National Health Service; EHR: Electronic Health Record; HL7: Health Level 7; SQL: Structured Query Language; OLAP: Online Analytical Processing; KCH: King’s College London; NLP: Natural Language Processing; OCR: Optical Character Recognition; IE: Information Extraction; SLAM: South London and Maudsley; NER: Named Entity Recognition; HPO: Human Phenotype Ontology; JSON: JavaScript Object Notation; PHI: Protected Health Identifiers; I2B2: The Informatics for Integrating Biology and the Bedside; JMS: Java Messaging Service; REST: Representational State Transfer; LDAP: Lightweight Directory Access Protocol; HTTP: Hypertext Transfer Protocol; SSL: Secure Socket Layer; 100KGP: 100 000 Genomes Project; CCP: Cyclic Citrullinated Peptide; API: Application Programming Interface; CRIS: Clinical Records Interactive Search; SAIL: Secure Anonymised Data Link.

### Declarations

#### Ethics approval and consent to participate

The creation of the CogStack software was an internal service development project for King’s College Hospital NHS Foundation Trust, and thus did not require ethical approval. As no patient identifiable data was required for the development of the software, no approval was sought from the Health Research Authority according to Confidentiality Advisory Group guidelines (http://www.hra.nhs.uk/resources/confidentiality-advisory-group/determining-need-cag-application/). The validation of the Bio-YODIE software made use of the CRIS dataset, which is approved as an anonymised data resource for secondary analysis by Oxfordshire Research Ethics Committee C (08/H0606/71) and governance is provided for all projects and dissemination through a patient-led oversight committee.

### Consent to publish

Not applicable: No individual persons data is presented in this manuscript.

### Availability of data and materials

Not applicable

### Funding

This paper represents independent research part funded by the National Institute for Health Research (NIHR) Biomedical Research Centre at South London and Maudsley NHS Foundation Trust and King’s College London, the University College London Hospitals Biomedical Research Centre, by awards establishing the Farr Institute of Health Informatics Research at UCL Partners, from the Medical Research Council, Arthritis Research UK, British Heart Foundation, Cancer Research UK, Chief Scientist Office, Economic and Social Research Council, Engineering and Physical Sciences Research Council, National Institute for Health Research, National Institute for Social Care and Health Research, and Wellcome Trust (grant MR/K006584/1). Additional funding came from the following awards. NHS England Enablement funding, the UK Infrastructure for Large-scale Clinical Genomics Research *MC*_*P*_ *C*_1_4089 and the European Union’s Horizon 2020 research and innovation programme under grant agreement No 644753 (KConnect). The views expressed are those of the author(s) and not necessarily those of the NHS, the NIHR or the Department of Health.

### Authors’ contributions

RJ led the architecture for CogStack, developed the batch process, data simulation algorithms, and wrote the manuscript. RJ and IK developed the business requirements for Cognition. IK led the design and development of Cognition, including the de-identification algorithms used here, and offered additional contributions to the manuscript. KL and AA provided additional development and technical support. DL, DN and CS provided project management and business support for KCH. TG provided support for Biolark. AR, GG, XS and HW provided support for Bio-YODIE. RS and RD provided supervision support.

## Acknowledgements

We would like to thank Dr Will Bernal, Caldicott Guardian for KCH for his advice on governance and ethical matters.

## Author details

^1^Institute of Psychiatry, Psychology and Neuroscience, King’s College London, 16 De Crespigne Park, London, UK SE5 8AF. ^2^South London and Maudsley NHS Foundation Trust, Denmark Hill, London, UK SE5 8AZ. ^3^King’s College Hospital, Denmark Hill, London, UK SE5 9RS. ^4^University of Sheffield, Western Bank, Sheffield, UK, S10 2TN. ^5^Farr Institute of Health Informatics Research, UCL Institute of Health Informatics, University College London, London, UK, WC1E 6BT. ^6^Garvan Institute of Medical Research, Sydney, Australia, NSW 2010.^7^InterDigital Communications, 64 Great Eastern Street, 1st Floor, London, UK, EC2A 3QR.

